# The effect of single-task training on learning transfer to a novel bimanual task

**DOI:** 10.1101/858217

**Authors:** Zaeem Hadi, Aqsa Shakeel, Hafsa Ahmad, Muhammad Nabeel Anwar, Muhammad Samran Navid

## Abstract

**Background:** The contextual interference effect suggests that the random practice of multiple-tasks is more beneficial for the retention and transfer of the learning as compared to blocked practice. Therefore, the transfer of learning is usually attributed to the contextual interference effect and is studied in a multi-task setting.

**Objective:** The goals of this study were to evaluate whether the transfer of learning (i) can occur when a single bimanual task is practiced, (ii) is affected by the knowledge of results (feedback), and (iii) sustains over an extended number of trials.

**Methods:** Fifty-two healthy subjects were equally divided into four groups. Before the transfer test, two groups practiced a bimanual finger-tapping task with feedback (EF) and without feedback (ENF). The third group (IM) practiced the same task using the kinesthetic motor imagery, whereas the last group acted as a control (CTRL) and performed only a bimanual button-pressing task used for the transfer test.

**Results:** Linear mixed-model showed that in the transfer test, groups EF, ENF and IM had similar performance but significantly higher scores compared to the CTRL group. Compared to the CTRL, the EF and IM groups showed significantly improved performance in most of the sessions but group ENF had similar results.

**Conclusion:** This study suggests that the single-task practice of a discrete bimanual task can facilitate the learning transfer to a novel task and knowledge of results (feedback) have no significant impact on the transfer of learning. Moreover, the transfer of learning effect does not disappear in extended trials.

**Highlights:**

- Single-task practice of a discrete bimanual task can facilitate the learning of a novel bimanual task
- Knowledge of results (feedback) does not improve learning transfer in single-task setting
- Transfer of learning effect does not disappear in extended trials

## 1 Introduction

The interference of practice tasks during multi-task learning is explained in terms of contextual interference. Contextual interference is the improvement in skill learning because of the interference during practice (Magill & Hall, 1990). The concept of contextual interference was first studied for motor skill acquisition in 1979 (Shea & Morgan, 1979). High contextual interference, which occurs during random practice, is associated with increased retention and learning transfer whereas low contextual interference, which takes place during blocked practice, is associated with the performance during acquisition (Shea & Morgan, 1979). The effects of contextual interference also hold for motor imagery tasks (Gabriele, Hall, & Lee, 1989).

Contextual interference studies employ motor tasks to evaluate motor skill learning. Motor tasks can either be unimanual or bimanual. Bimanual motor tasks such as driving or typing on a keyboard are more complex than unimanual motor tasks and require a high level of coordination (Yeganeh Doost, Orban de Xivry, Bihin, & Vandermeeren, 2017). In general, the coordinated in-phase and anti-phase motion of both limbs is preferred in bimanual tasks (Swinnen, Dounskaia, Walter, & Serrien, 1997). Non-iso frequencies (e.g. in the ratio of 1:2, 2:1 or 1:3) and polyrhythms (e.g. in the ratio of 3:2, 4:3 or 5:2) are difficult to perform (Peper, Beek, & van Wieringen, 1995) and often require feedback to be performed efficiently (Kovacs & Shea, 2011). Most of the contextual interference studies that looked at the learning of bimanual patterns have yielded contradictory results and the beneficial effect of random practice on retention and transfer was either not observed or observed partly (Maslovat, Chua, Lee, & Franks, 2004; Rey, Liu, & Simpson, 1994; Tsutsui, Lee, & Hodges, 1998). Studies have commonly employed continuous bimanual patterns, such as pattern making and tracking (Maslovat et al., 2004; Pauwels, Swinnen, & Beets, 2014; Tsutsui et al., 1998), or sequential bimanual tasks, such as sequential key pressing (Rey et al., 1994). To the best of our knowledge, the discrete bimanual tasks, such as finger tapping or button pressing, are not used in contextual interference studies.

Contextual interference can be affected by the knowledge of results (KR) or feedback (Del Rey & Shewokis, 1993; Patterson, Carter, & Hansen, 2013; Russell & Newell, 2007), which is often ignored in the studies. KR plays an important role in the learning of motor skills but its effects on contextual interference are not well documented. Some studies have proposed that KR affects the retention and transfer of learning (Del Rey & Shewokis, 1993; Patterson et al., 2013; Russell & Newell, 2007) whereas (Wu et al., 2011) reported that KR effect is more likely to be overshadowed by the type of practice schedule.

Previous studies have established a reasonable understanding of contextual interference effect (Christina, 2017; Gabriele et al., 1989; Pauwels et al., 2014; Shea & Morgan, 1979) but a number of studies have reported contradictory results (Buszard, Reid, Krause, Kovalchik, & Farrow, 2017; Caramiaux, Bevilacqua, Wanderley, & Palmer, 2018; Moretto, Marcori, & Okazaki, 2018; Russell & Newell, 2007). The contextual interference effect requires the practice of multiple variations of a task (Magill & Hall, 1990) and thus, the retention and transfer of learning effects were significantly higher in multi-task practice as compared to single-task practice (Maslovat et al., 2004). VaezMousavi et al. (VaezMousavi & Rostami, 2009) reported that transfer of learning can occur when learning a single-task (basketball throw), however, it has also been shown in multi-task practice that transfer of learning tends to disappear when tested over multiple sessions (also called extended or delayed trials) (Meira & Tani, 2003; Meira Jr & Tani, 2001; Perez, Meira Jr, & Tani, 2005).

From the previous studies, it is evident that the transfer of learning is not usually tested for single-task practice and the effect of extended trials on the transfer of learning remains doubtful when practicing a single-task. Moreover, the impact of KR (feedback) is largely ignored, and bimanual tasks have mostly yielded contradictory results in such studies. Thus, the objectives of the present study were to evaluate whether (i) single-task practice of a discrete-bimanual task can facilitate the transfer of learning, (ii) KR (feedback) affects the transfer of learning, and (iii) transfer of learning decrease in extended trials. For the first objective, we utilized a finger-tapping task for practice and button-pressing task for the transfer of learning test, whereas for the second objective, one of the groups performed the button-pressing task (test session) with KR (feedback). The third objective was evaluated using multiple sessions for the transfer test.

## 2 Methods

The protocol of the study was approved by the local ethics committee of the National University of Sciences and Technology (NUST), Islamabad, Pakistan, in accordance with the Declaration of Helsinki.

### 2.1 Subjects

Fifty-two right-handed healthy university students took part in this study. All subjects included in the study were naive to similar studies and tasks used in the experiment. This was done to remove any bias or individual differences. Before participation, all subjects signed written informed consent. Subjects were instructed to have a good sleep and not take tea or coffee before the experiment. Verbal and written instructions about the tasks were provided to each subject.

The subjects were randomly divided into four groups (n = 13/group), execution with feedback (EF) (age: 25.6 ± 1.2 years, 8 males), execution without feedback (ENF) (age: 24.0 ± 1.2 years, 6 males), imagery (IM) (age: 22.9 ± 2.8 years, 8 males) and control (CTRL) (age: 24.3 ± 2.1 years, 7 males). The EF, ENF and IM groups performed both training and testing tasks, whereas the CTRL group only performed the testing task.

### 2.2 Motor imagery ability

Before the start of the experiment, the IM group filled the Kinesthetic and Visual Imagery (KVIQ-10) questionnaire (Malouin et al., 2007). The purpose of the questionnaire was to get subjects acquainted with the type of imagery they had to perform. KVIQ-10 questionnaire contained 5 questions for kinesthetic imagery ability. The visual imagery part of the questionnaire was not administered as subjects were instructed to perform only kinesthetic imagery. The subjects were first verbally explained the type of movement to be performed and then demonstrated by the experimenter. Afterward, subjects were instructed to perform the same movement and then imagine the proprioceptive sensation of the movement (kinesthetic imagery; KI). The kinesthetic imagery score was measured on a scale from 1 to 5, where 1 on the scale was equivalent to ‘no sensation’ and 5 was equivalent to ‘as intense as executing the action’. The total score of the questionnaire ranged from 5 to 25. The mean kinesthetic imagery score of the IM group was 13.92 ± 4.13.

### 2.3 Protocol

The experiment was composed of a practice session and a testing session, separated by 15 min. A bimanual finger-tapping task was utilized in the practice session, whereas a bimanual button-pressing task was used for the transfer test session. Auditory stimuli were provided to mark the start of tasks (finger tap/button press) through earphones. The feedback in the EF group and the stimuli were generated using MATLAB 2012a (MathWorks®, Inc., Natick, MA, USA.) and Psychtoolbox-3 (PTB-3) (Brainard & Vision, 1997). The required movements were recorded at a sampling rate of 1000 Hz using a pulse transducer (MLT1010/D, AD Instruments, Australia), a push button (MLA92/D, AD Instruments, Australia) and two custom made resistive touch sensors, connected to PowerLab 26T (AD Instruments, Australia). The data was recorded in real-time using LabChart (v7.3.7, AD Instruments, Australia). The number of correct trials in the transfer test was scored manually by visual inspection of the data in LabChart.

### 2.4 Tasks

#### 2.4.1 Training

The training sessions were performed by the EF, ENF and IM groups. For training, the finger-tapping task was adopted from a previous study (Shakeel, Ahmad, Navid, Mahroo, & Anwar, 2017). Subjects sat on a chair 0.6 m away from a computer screen. Subjects were required to perform bimanual finger tapping in 2:1 frequency with index fingers on a resistive touch sensor fixed on the table. The 2:1 tapping frequency specified simultaneous tapping of index fingers of both hands for one tap, followed by tapping of the index finger of the non-dominant (left) hand. The dominant (right) index finger was held static at peak upward position during left index finger tapping. The left side of the diagonal line in Figure 1 gives an overview of the training session.

**Figure 1.**
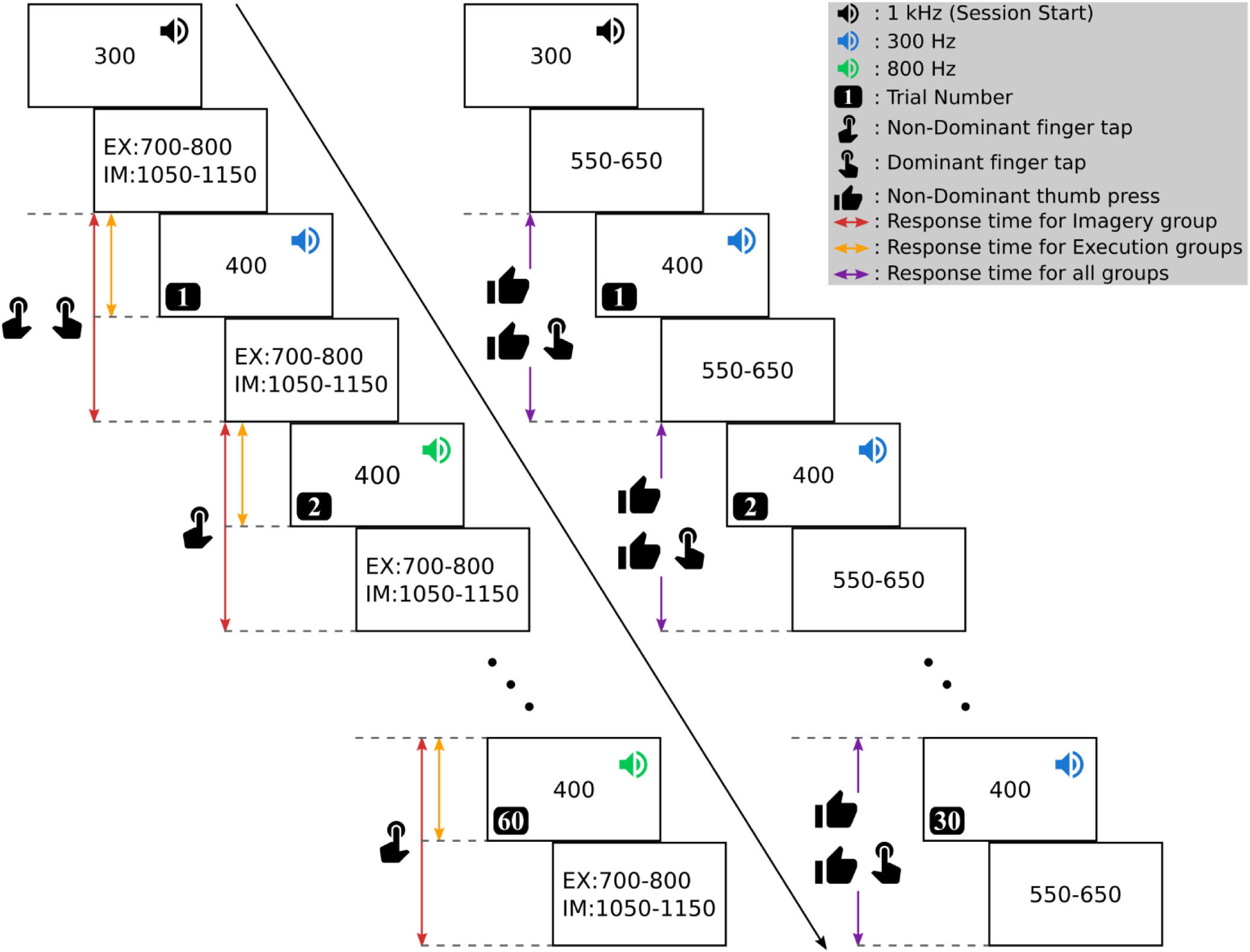
Methodology overview. One session of training (left side of the time direction diagonal) and one session of testing (right side of the time direction diagonal) is shown. The value in the center of each rectangular box is time in ms. The movements shown horizontally together were required to be performed in synchronization. Abbreviations: EX = Execution groups, IM = Imagery group.

The training had 14 sessions with 60 trials per session. There was a 1-minute break between each session. A 1 kHz auditory stimulus for 300 ms was presented that marked the start of every session. Afterward, 400ms long low frequency (300 Hz) and high frequency (800 Hz) auditory stimuli were given alternatively. The time between any two stimuli was 700-800 ms for the EF and ENF groups, and 1050-1150 ms for the IM group. The interval was kept larger for IM group than the execution groups to ensure that the subjects completed each imagery trial, as there was no data being recorded. The subjects were required to execute/imagine the tapping of index fingers of both hands on the low frequency stimulus, and tapping of index finger of non-dominant hand on the high frequency stimulus.

The ENF group was instructed to perform the tapping as soon as they receive the stimulus, whereas the EF group was required to perform the tapping within the stimulus duration. The EF group received visual feedback on the screen about the response time by a colored bar (a green bar indicated accurate execution and a red bar represented error). A trial was considered correct if the tap was performed before the completion of 400 ms stimulus duration. The IM group was asked to perform the imagery of the tapping before the presentation of next auditory stimulus.

#### 2.4.2 Transfer test

The testing sessions were performed by all groups. For the transfer test, the subjects were required to perform tapping with the index finger of the dominant hand on a pulse transducer fixed on the table along with pressing the push-button twice with the thumb of the non-dominant hand instantly after hearing the auditory stimulus. The subjects were instructed to synchronize their tap with the second press of the button. The right side of the diagonal line in Figure 1 gives an overview of a testing session.

The testing was comprised of 7 sessions with 30 trials per session. There was a 1-minute break between sessions. An auditory stimulus of 1 kHz frequency with a duration of 300 ms marked the start of the session. Afterward, 400 ms wide 300 Hz auditory stimuli were generated. The duration between consecutive stimuli was 550-650 ms. The subjects were instructed to execute the movements before the generation of the next auditory stimulus. A trial was considered correct if the movement was completed before the next auditory stimulus.

### 2.5 Statistical Analysis

R version 3.5.1 (Team, 2018) was used to perform statistical analysis. The data are presented as mean ± SD unless otherwise indicated. The statistical significance threshold was set at *p* < 0.05.

Linear mixed-effect model (LMM) was used to identify whether (a) the groups are different based on the scores obtained in the transfer test (objectives (i) and (ii)), and (b) the scores are different across sessions (objective (iii)). The between-subject variance was estimated using a random intercept in the model. The model was implemented using ‘lme4’ package (Bates, Mächler, Bolker, & Walker, 2015) version 1.1.18.1 in R using the syntax:

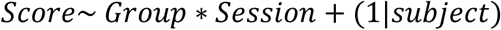

We evaluated the fit of the model using Akaike information criterion corrected (AICc) for small samples, where models were made using gaussian, gamma and poisson distributions. The model with gaussian distribution had the lowest AICc and the most well fitted best-fit line in the fitted vs residual plot, therefore, it was used for further analysis. The package ‘emmeans’ (Lenth, 2018) version 1.2.4 was used for obtaining the contrasts adjusted for multiple comparisons using Tukey’s test.

## 3 Results

All subjects successfully completed the training and transfer test sessions, and data from all of them were included for analysis. The scores for each subject across all transfer test sessions are plotted in the Figure 2. There was more variance in the score of CTRL and IM groups compared to EF and ENF group, as can be seen by the distribution of the boxplots.

**Figure 2.**
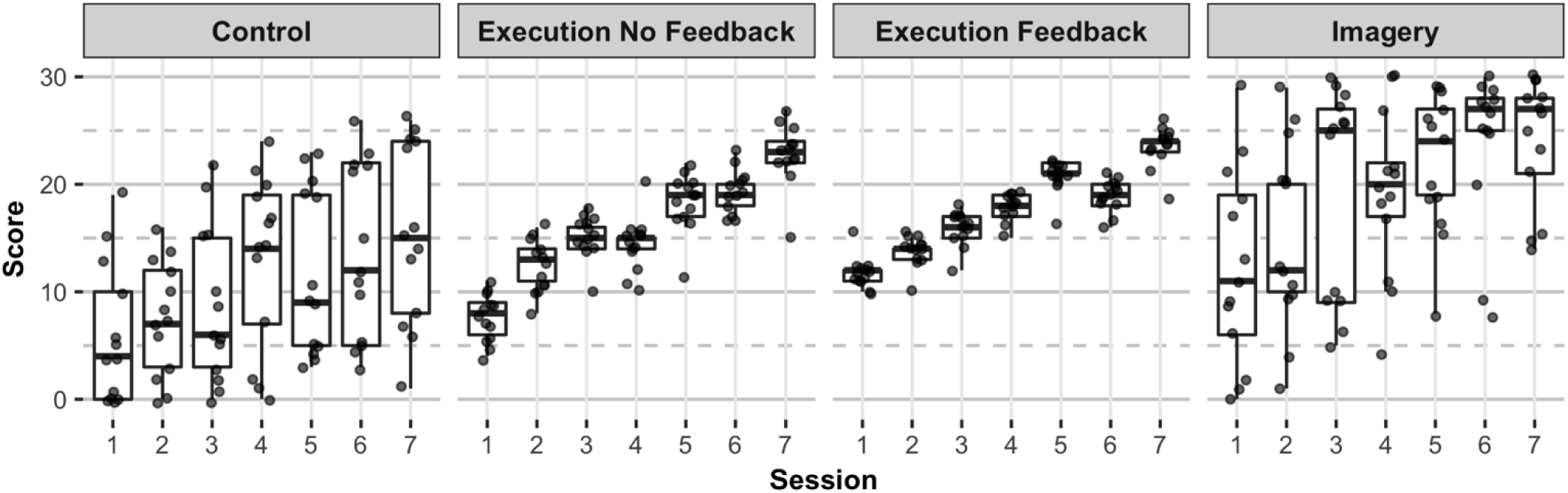
Distribution of raw scores. Boxplots show the median, 25th and 75th percentiles. The data show a positive trend suggesting increase in the correct responses, however, there is more variance in score of control and imagery groups.

The LMM showed significant effects of both factors; group (χ^2^(3) = 29.39, *p* < 0.001) and session (χ^2^(6) = 407.96, *p* < 0.001).

Table 1 and Table 2 show the effect sizes from the model for groups and sessions, respectively. The pairwise contrasts are given in Table 3 for groups and show that the groups EF, ENF and IM have significantly higher mean scores from the CTRL group, however, there is no difference among themselves. The pairwise contrasts are given in Table 4 for sessions and show that the scores on average increased in each session compared to the previous sessions except for the 4^th^, 5^th^ and 6^th^ session (compared to their immediate previous session). Figure 3 shows the pairwise contrasts for group and session.

**Table 1.** Estimated effect size for the factor group from the statistical model. Results are averaged over the levels of session. Significant effects (*p*<0.05) are in bold text.

| Group | Estimated Mean Score ± SE | 95% CI | *t* ratio | *p* value |
| --- | --- | --- | --- | --- |
| Control | 10.89 ± 1.22 | [8.44, 13.34] | 8.92 | < **.0001** |
| Execution No Feedback | 15.71 ± 1.22 | [13.27, 18.16] | 12.87 | < **.0001** |
| Execution Feedback | 17.47 ± 1.22 | [15.03, 19.92] | 14.31 | < **.0001** |
| Imagery | 19.48 ± 1.22 | [17.04, 21.93] | 15.96 | < **.0001** |

**Table 2.** Estimated effect size for the factor session from the statistical model. Results are averaged over the levels of group. Significant effects (*p*<0.05) are in bold text.

| Session | Estimated Mean Score ± SE | 95% CI | *t* ratio | *p* value |
| --- | --- | --- | --- | --- |
| 1 | 9.38 ± 0.78 | [7.84, 10.93] | 12.00 | < **.0001** |
| 2 | 12.25 ± 0.78 | [10.70, 13.80] | 15.66 | < **.0001** |
| 3 | 14.85 ± 0.78 | [13.30, 16.39] | 18.98 | < **.0001** |
| 4 | 16.08 ± 0.78 | [14.53, 17.62] | 20.55 | < **.0001** |
| 5 | 18.21 ± 0.78 | [16.67, 19.76] | 23.28 | < **.0001** |
| 6 | 19.00 ± 0.78 | [17.45, 20.55] | 24.29 | < **.0001** |
| 7 | 21.46 ± 0.78 | [19.92, 23.01] | 27.44 | < **.0001** |

**Table 3.** Estimated contrasts for the factor group from the statistical model. Results are averaged over the levels of session. Significant effects (*p*<0.05) are in bold text.

| Group | Difference ± SE | 95% CI | *t* ratio | *p* value |
| --- | --- | --- | --- | --- |
| Control - Execution No Feedback | -4.82 ± 1.73 | [-9.40, -0.25] | -2.79 | **0.0348** |
| Control - Execution Feedback | -6.58 ± 1.73 | [-11.15, -2.01] | -3.81 | **0.0019** |
| Control - Imagery | -8.59 ± 1.73 | [-13.16, -4.02] | -4.98 | **< .0001** |
| Execution No Feedback - Execution Feedback | -1.76 ± 1.73 | [-6.33, 2.81] | -1.02 | 0.7395 |
| Execution No Feedback - Imagery | -3.77 ± 1.73 | [-8.34, 0.80] | -2.18 | 0.1405 |
| Execution Feedback - Imagery | -2.01 ± 1.73 | [-6.58, 2.56] | -1.17 | 0.6514 |

**Table 4.**
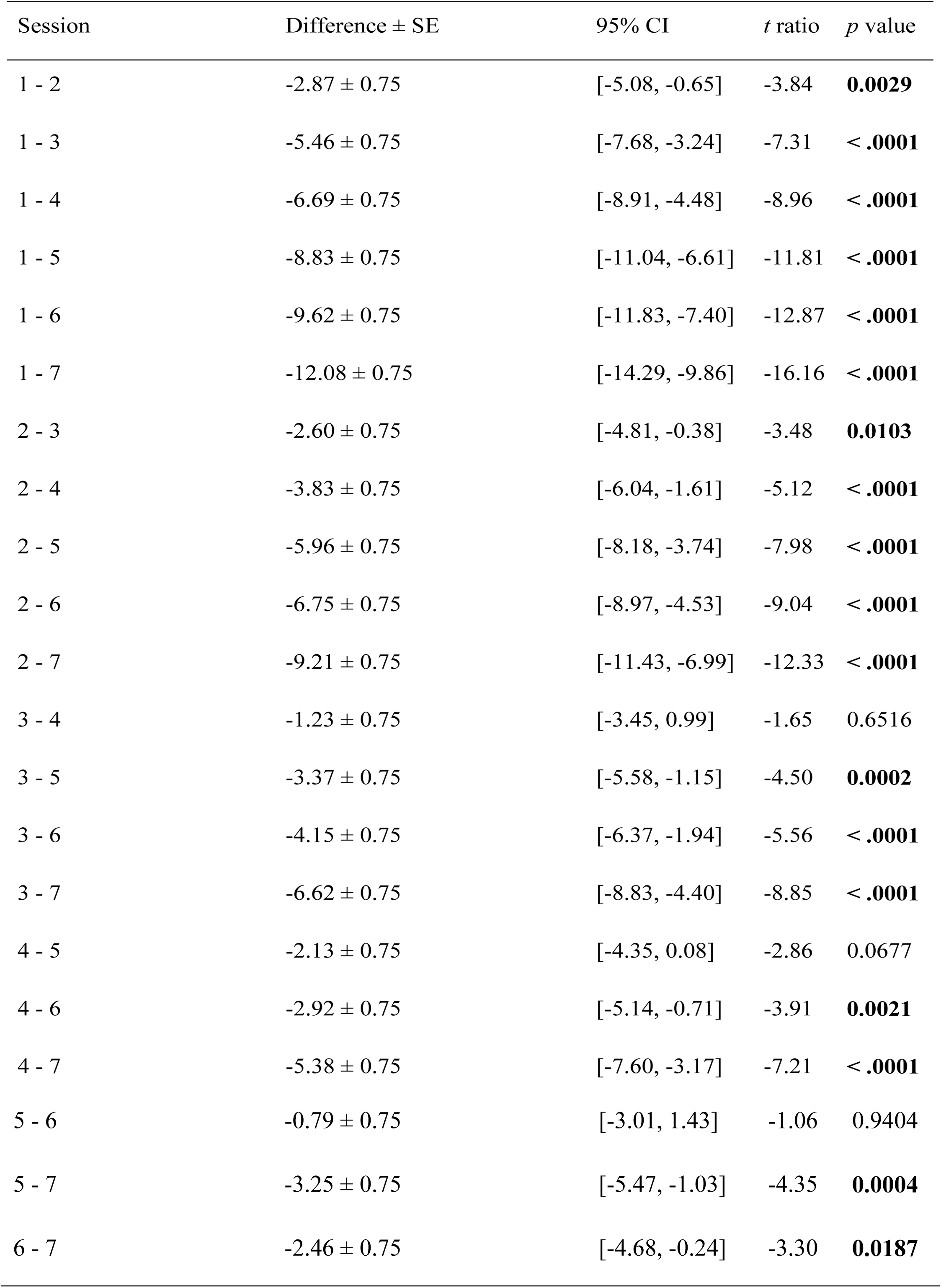
Estimated contrasts for the factor session from the statistical model. Results are averaged over the levels of group. Significant effects (*p*<0.05) are in bold text.

**Figure 3.**
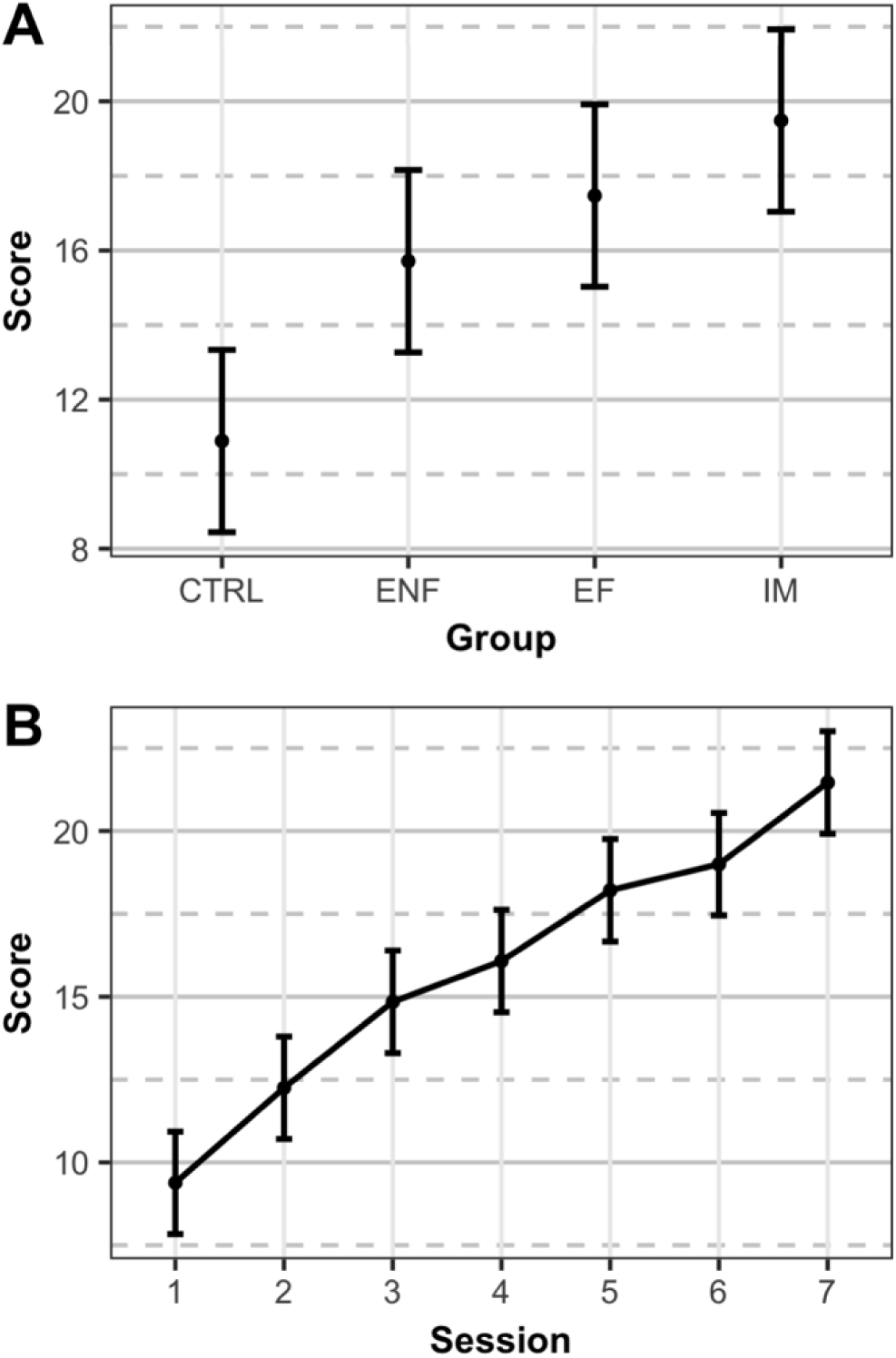
Pairwise contrasts. Error bar shows estimated mean score ± 95% CI. (A) The scores across groups (averaged over sessions) showed significant difference between the CTRL and other groups, whereas there were no significant differences among the EF, ENF and IM groups. (B) The scores increased (averaged over groups) significantly from the first session to the last session, except for sessions 4-6 which were not significantly different from their immediate previous session. Abbreviations: (group) CTRL = Control, ENF = Execute without feedback, EF = Execute with feedback, IM = Imagery.

Table 5 shows the pairwise comparisons of the groups over each session. By comparing the practice groups EF, ENF, and IM to the CTRL group to test whether the transfer of learning disappears over extended trials, it was found that the practice groups had significant differences over some sessions. There were no differences among the groups EF, ENF, and IM. The ENF group did not show any improved performance compared to the CTRL group in sessions 1, 2, 4, and 6. The performance of group EF was significantly better in most of the sessions whereas the IM group was significantly better than the CTRL group in all sessions.

**Table 5.** Estimated contrasts for interaction of session and group factors from the statistical model. Significant effects (p<0.05) are in bold text.

| Contrast | Session | Difference ± SE | 95% CI | *t* ratio | *p* value |
| --- | --- | --- | --- | --- | --- |
| Control - Execution No Feedback | 1 | -1.69 ± 2.20 | [-7.42, 4.04] | -0.768 | 0.8687 |
| Control - Execution Feedback | 1 | -5.77 ± 2.20 | [-11.50, -0.04] | -2.619 | **0.0477** |
| Control - Imagery | 1 | -6.38 ± 2.20 | [-12.11, -0.66] | -2.898 | **0.0224** |
| Execution No Feedback - Execution Feedback | 1 | -4.08 ± 2.20 | [-9.81, 1.65] | -1.851 | 0.2544 |
| Execution No Feedback - Imagery | 1 | -4.69 ± 2.20 | [-10.42, 1.04] | -2.130 | 0.1487 |
| Execution Feedback - Imagery | 1 | -0.62 ± 2.20 | [-6.34, 5.11] | -0.279 | 0.9924 |
| Control - Execution No Feedback | 2 | -4.84 ± 2.20 | [-10.57, 0.88] | -2.200 | 0.1284 |
| Control - Execution Feedback | 2 | -6.23 ± 2.20 | [-11.96, -0.50] | -2.828 | **0.0272** |
| Control - Imagery | 2 | -7.77 ± 2.20 | [-13.50, -2.04] | -3.526 | **0.0032** |
| Execution No Feedback - Execution Feedback | 2 | -1.38 ± 2.20 | [-7.11, 4.34] | -0.628 | 0.9227 |
| Execution No Feedback - Imagery | 2 | -2.92 ± 2.20 | [-8.65, 2.81] | -1.327 | 0.5475 |
| Execution Feedback - Imagery | 2 | -1.54 ± 2.20 | [-7.27, 4.19] | -0.698 | 0.8976 |
| Control - Execution No Feedback | 3 | $-6.38 \pm 2.20$ | [-12.11, -0.66] | -2.898 | **0.0224** |
| Control - Execution Feedback | 3 | $-7.08 \pm 2.20$ | [-12.81, -1.35] | -3.212 | **0.0088** |
| Control - Imagery | 3 | $-10.85 \pm 2.20$ | [-16.57, -5.12] | -4.923 | **< .0001** |
| Execution No Feedback - Execution Feedback | 3 | $-0.69 \pm 2.20$ | [-6.42, 5.04] | -0.314 | 0.9892 |
| Execution No Feedback - Imagery | 3 | $-4.46 \pm 2.20$ | [-10.19, 1.27] | -2.025 | 0.1837 |
| Execution Feedback - Imagery | 3 | $-3.77 \pm 2.20$ | [-9.50, 1.96] | -1.711 | 0.3219 |
| Control - Execution No Feedback | 4 | $-1.46 \pm 2.20$ | [-7.19, 4.27] | -0.663 | 0.9106 |
| Control - Execution Feedback | 4 | $-4.85 \pm 2.20$ | [-10.57, 0.88] | -2.200 | 0.1284 |
| Control - Imagery | 4 | $-6.30 \pm 2.20$ | [-12.04, -0.58] | -2.863 | **0.0247** |
| Execution No Feedback - Execution Feedback | 4 | $-3.38 \pm 2.20$ | [-9.11, 2.34] | -1.536 | 0.4187 |
| Execution No Feedback - Imagery | 4 | $-4.85 \pm 2.20$ | [-10.57, -0.88] | -2.200 | 0.1284 |
| Execution Feedback - Imagery | 4 | $-1.46 \pm 2.20$ | [-7.19, 4.27] | -0.663 | 0.9106 |
| Control - Execution No Feedback | 5 | $-6.54 \pm 2.20$ | [-12.27, -0.81] | -2.968 | **0.0183** |
| Control - Execution Feedback | 5 | $-9.00 \pm 2.20$ | [-14.73, -3.27] | -4.085 | **0.0004** |
| Control - Imagery | 5 | -10.23 ± 2.20 | [-15.96, -4.50] | -4.644 | **< .0001** |
| Execution No Feedback - Execution Feedback | 5 | -2.46 ± 2.20 | [-8.19, 3.27] | -1.117 | 0.6795 |
| Execution No Feedback - Imagery | 5 | -3.69 ± 2.20 | [-9.42, 2.04] | -1.676 | 0.3403 |
| Execution Feedback - Imagery | 5 | -1.23 ± 2.20 | [-6.96, 4.50] | -0.559 | 0.9440 |
| Control - Execution No Feedback | 6 | -5.62 ± 2.20 | [-11.34, 0.11] | -2.549 | 0.0569 |
| Control - Execution Feedback | 6 | -5.23 ± 2.20 | [-10.96, 0.49] | -2.374 | 0.0868 |
| Control - Imagery | 6 | -10.08 ± 2.20 | [-15.81, -4.35] | -4.574 | **0.0001** |
| Execution No Feedback - Execution Feedback | 6 | 0.38 ± 2.20 | [-5.34, 6.11] | 0.175 | 0.9981 |
| Execution No Feedback - Imagery | 6 | -4.46 ± 2.20 | [-10.19, 1.27] | -2.025 | 0.1837 |
| Execution Feedback - Imagery | 6 | -4.85 ± 2.20 | [-10.57, 0.88] | -2.200 | 0.1284 |
| Control - Execution No Feedback | 7 | -7.23 ± 2.20 | [-12.96, -1.50] | -3.282 | **0.0070** |
| Control - Execution Feedback | 7 | -7.92 ± 2.20 | [-13.65, -2.19] | -3.596 | **0.0025** |
| Control - Imagery | 7 | -8.54 ± 2.20 | [-14.27, -2.81] | -3.876 | **0.0009** |
| Execution No Feedback - Execution Feedback | 7 | -0.69 ± 2.20 | [-6.42, 5.04] | -0.314 | 0.9892 |
| Execution No Feedback -<br>Imagery | 7 | -1.31 ± 2.20 | [-7.04, 4.42] | -0.594 | 0.9339 |
| Execution Feedback -<br>Imagery | 7 | -0.62 ± 2.20 | [-6.34, 5.11] | -0.279 | 0.9924 |

## 4 Discussion

This study was designed to develop an understanding of the transfer of learning effect in a single-task learning setup by primarily focusing on learning of discrete-bimanual patterns. Moreover, we examined whether KR (feedback) affects transfer of learning, and if the transfer of learning effect disappears in extended trials during single-task practice. Our results showed that single-task practice facilitated the transfer of learning in discrete-bimanual task. All training groups EF, ENF and IM performed significantly better in the transfer test as compared to the CTRL group. However, the feedback did not significantly affect the transfer of learning, since EF, ENF and IM groups showed similar scores. On the other hand, the effect of transfer of learning did not disappear over time.

Learning of finger-tapping task facilitated the performance in transfer test using button-pressing task. Our results are comparable to a previous study, which used a basketball free throw as the single practice task (VaezMousavi & Rostami, 2009), thus suggesting that transfer of learning effect in applied settings can also be generalized to laboratory settings during single task practice.

The IM group had the highest mean score but was statistically similar to the groups EF and ENF. However, it was significantly better than the CTRL group. These results are also comparable to the findings of (VaezMousavi & Rostami, 2009), in which the imagery group achieved the highest score in comparison to all other groups in retention- and transfer-tests. A possible explanation of the highest mean score of IM group could be the spacing effect (i.e. difference in trial duration between groups) explained in (Richland, Finley, & Bjork, 2004) where random practice group with longer trial duration performed better in transfer test as compared to the random practice group with shorter trial duration. In our study, the trial duration of the IM group was longer as compared to those of the practice groups EF and ENF. Thus, longer duration to respond may have played a role in the IM group obtaining the highest mean score, but this needs to be further investigated. The higher variability in the performance of the IM group can also be a reason for achieving the highest mean score. Thus, we would like to suggest that motor imagery practice during acquisition is likely able to produce superior or similar performance in transfer test as compared to the physical practice, but at the expense of higher variability.

We evaluated if KR (feedback) results in superior performance in the transfer test. We found that the EF group outperformed the CTRL group only. These results are in contrast to the previous studies such as (Del Rey & Shewokis, 1993; Patterson et al., 2013; Russell & Newell, 2007). Del Rey et al. (Del Rey & Shewokis, 1993) compared the difference of performance in case of full KR (feedback after every trial) and spaced KR (feedback after every 5 or 10 trials), but in our study, we compared the effect of KR or no KR using separate groups (EF and ENF). In Russel et al. (Russell & Newell, 2007), retention tests also had KR (feedback). This is important to mention, as feedback is not often provided in retention and transfer tests. Patterson et al. (Patterson et al., 2013) did find an effect of KR but the presence and absence of KR were controlled by the subjects. Thus, not having a universally controlled factor could have been a source of bias in the results. A 100% KR (feedback after each trial) is considered to deteriorate retention and transfer performance (Wu et al., 2011), however, no such effect was observed in our study and there was no difference between groups EF and ENF. The discrete bimanual pattern of 2:1 is considered to make bimanual performance more challenging compared to its reverse arrangement (Semjen, 2002) and it is suggested that complex modes require feedback to be efficiently performed and its absence would decline the performance (Kovacs & Shea, 2011). However, this was not clearly observable in our study. Although, the group EF performed slightly better than group ENF in all sessions except in session 6, however, the difference is negligible. Thus, we would like to suggest that KR may slightly enhance transfer test performance, but the improvement may not be statistically significant.

This study also aimed to establish a relationship between the effects of learning transfer and the discrete bimanual tasks. From the previous studies that have used continuous bimanual tasks (Maslovat et al., 2004; Pauwels et al., 2014; Tsutsui et al., 1998) and sequential bimanual tasks (Rey et al., 1994), some have yielded results that did not support the retention and transfer of learning effects (Maslovat et al., 2004; Rey et al., 1994; Tsutsui et al., 1998). Our study suggests that transfer of learning can occur in discrete-bimanual tasks such as finger-tapping or button-pressing. However, we used a single task practice design. Further investigation is required, especially in bimanual task-based studies using either single-task or multi-task setting, to generalize the results.

From previous studies, it was expected that the transfer of learning effect dissipates over extended trials (Meira & Tani, 2003; Meira Jr & Tani, 2001; Perez et al., 2005). The scores in groups EF and IM were significantly better than the CTRL group and the transfer of learning effect did not disappear in extended trials. The scores of the ENF group were overall comparable to the CTRL group and, therefore, the effect of extended trials was not observed. The improvement in the performance of groups EF, ENF, and IM compared to the CTRL group cannot be solely attributed to motor learning because the CTRL group performed an equal number of sessions and trials of the same task. Previous studies (Meira & Tani, 2003; Meira Jr & Tani, 2001; Perez et al., 2005) refer to similar effects when extended or delayed trials are discussed but their measure for the consideration of transfer test trials as extended trials is different. Meira et al. (Meira & Tani, 2003) performed the first transfer test 24 hours after completing the practice and the second transfer test 24 hours after completing first transfer test, and this was considered delayed transfer test (comparable to extended trials). Perez et al. (Perez et al., 2005) also used two transfer tests with a difference of one day between them. However, Meira Jr et al. (Meira Jr & Tani, 2001) considered every increasing block (session) of transfer test trials to be an extended trial. Our study design is comparable to Meira Jr et al. (Meira Jr & Tani, 2001) in the sense that we also considered each increasing block (session) of transfer test trials to be extended trials, however, we found no relation of extended trials with the transfer of learning. Thus, we suggest that the effect of extended trials on the transfer of learning is only observable in a multi-task setting as in the contextual interference effect and not in a single-task setting.

## 5 Limitations

We trained subjects using the kinesthetic part of the KVIQ-10 questionnaire, but no empirical measure of motor imagery ability or motor imagery practice was used in the analysis. Furthermore, we increased the trial duration of motor imagery practice trials to ensure that subjects have enough time to complete the trial. To overcome these limitations, EEG can be used as an empirical measure, which can also explain the temporal dynamics of motor imagery leading to a better estimation of the trial duration. Efforts were made to compensate for individual differences as we recruited subjects who were naive to such studies and experimental tasks. Additionally, it was not possible to train the subjects to asymptotic levels before the experiment to remove any bias since the training conditions were different. Finally, the power of the study can be increased by increasing the sample size.

## 6 Conclusion

We conclude that single-task practice of a discrete bimanual task can facilitate the learning transfer to a novel task in a laboratory setting. We suggest that motor imagery practice during acquisition can have similar or better performance in transfer test as compared to the physical practice. Furthermore, we found that presence of KR and the extended trials have no impact on learning transfer when practicing a single-task.

## Author Contributions

Conceptualization: AS, MNA; Data curation: ZH, AS; Formal Analysis: ZH, MSN; Funding acquisition: MNA; Investigation: ZH, AS; Methodology: ZH, AS, MSN; Project administration: ZH, AS, MNA; Resources: MNA; Software: ZH, AS, HA, MSN; Supervision: MNA; Validation: ZH, MSN; Visualization: MSN; Writing – original draft: ZH, AS; Writing – review & editing: ZH, AS, HA, MNA, MSN.

## References

Bates, Douglas, Mächler, Martin, Bolker, Ben, & Walker, Steve. (2015). Fitting Linear Mixed-Effects Models Using lme4. Journal of Statistical Software, 67(1), 1–48.

Brainard, David H, & Vision, Spatial. (1997). The psychophysics toolbox. Spatial vision, 10, 433–436.

Buszard, Tim, Reid, Machar, Krause, Lyndon, Kovalchik, Stephanie, & Farrow, Damian. (2017). Quantifying Contextual Interference and Its Effect on Skill Transfer in Skilled Youth Tennis Players. Frontiers in psychology, 8, 1931.

Caramiaux, Baptiste, Bevilacqua Frédéric, Wanderley, Marcelo M, & Palmer, Caroline. (2018). Dissociable effects of practice variability on learning motor and timing skills. PloS one, 13(3), e0193580.

Christina, Robert W. (2017). Motor control and learning in the North American Society for the Psychology of Sport and Physical Activity (NASPSPA): The first 40 years. Kinesiology Review, 6(3), 221–231.

Del Rey, Patricia, & Shewokis, Patricia. (1993). Appropriate summary KR for learning timing tasks under conditions of high and low contextual interference. Acta psychologica, 83(1), 1–12.

Gabriele, Tina E, Hall, Craig R, & Lee, Timothy D. (1989). Cognition in motor learning: Imagery effects on contextual interference. Human Movement Science, 8(3), 227–245.

Kovacs, Attila J, & Shea, Charles H. (2011). The learning of 90 continuous relative phase with and without Lissajous feedback: external and internally generated bimanual coordination. Acta psychologica, 136(3), 311–320.

Lenth, Russell. (2018). emmeans: Estimated Marginal Means, aka Least-Squares Means.

Magill, Richard A, & Hall, Kellie G. (1990). A review of the contextual interference effect in motor skill acquisition. Human movement science, 9(3-5), 241–289.

Malouin, Francine, Richards, Carol L, Jackson, Philip L, Lafleur, Martin F, Durand, Anne, & Doyon, Julien. (2007). The Kinesthetic and Visual Imagery Questionnaire (KVIQ) for assessing motor imagery in persons with physical disabilities: a reliability and construct validity study. Journal of Neurologic Physical Therapy, 31(1), 20–29.

Maslovat, Dana, Chua, Romeo, Lee, Timothy D, & Franks, Ian M. (2004). Contextual interference: single task versus multi-task learning. Motor control, 8(2), 213–233.

Meira, CM, & Tani, G. (2003). Contextual interference effects assessed by extended transfer trials in the acquisition of the volleyball serve. Journal of Human movement studies, 45(5), 449–468.

Meira Jr, Cassio M, & Tani, Go. (2001). The contextual interference effect in acquisition of dart-throwing skill tested on a transfer test with extended trials. Perceptual and motor skills, 92(3), 910–918.

Moretto, Nelson Alexandre, Marcori, Alexandre Jehan, & Okazaki, Victor Hugo Alves. (2018). Contextual interference effects on motor skill acquisition, retention and transfer in sport rifle shooting. Human Movement, 19(2), 99–104.

Patterson, Jae T., Carter, Michael J., & Hansen, Steve. (2013). Self-controlled KR schedules: Does repetition order matter? Human Movement Science, 32(4), 567–579. doi: https://doi.org/10.1016/j.humov.2013.03.005

Pauwels, Lisa, Swinnen, Stephan P, & Beets, Iseult AM. (2014). Contextual interference in complex bimanual skill learning leads to better skill persistence. PloS one, 9(6), e100906.

Peper, CE, Beek, Peter J, & van Wieringen, Piet CW. (1995). Multifrequency coordination in bimanual tapping: Asymmetrical coupling and signs of supercriticality. Journal of Experimental Psychology: Human Perception and Performance, 21(5), 1117.

Perez, Carlos Rey, Meira Jr, Cassio M, & Tani, Go. (2005). Does the contextual interference effect last over extended transfer trials? Perceptual and Motor skills, 100(1), 58–60.

Rey, Patricia Del, Liu, Xiaoying, & Simpson, Kathy Jean. (1994). Does retroactive inhibition influence contextual interference effects? Research Quarterly for Exercise and Sport, 65(2), 120–126.

Richland, Lindsey E, Finley, Jason R, & Bjork, Robert A. (2004). Differentiating the contextual interference effect from the spacing effect. Paper presented at the Cognitive Science, Chicago.

Russell, Daniel M, & Newell, Karl M. (2007). How persistent and general is the contextual interference effect? Research quarterly for exercise and sport, 78(4), 318–327.

Semjen, Andras. (2002). On the timing basis of bimanual coordination in discrete and continuous tasks. Brain and Cognition, 48(1), 133–148.

Shakeel, Aqsa, Ahmad, Hafsah, Navid, Muhammad Samran, Mahroo, Amnah, & Anwar, Muhammad Nabeel. (2017). Performance feedback assists practice driven plasticity. Paper presented at the Biomedical Engineering (BioMed), 2017 13th IASTED International Conference on.

Shea, John B, & Morgan, Robyn L. (1979). Contextual interference effects on the acquisition, retention, and transfer of a motor skill. Journal of Experimental psychology: Human Learning and memory, 5(2), 179.

Swinnen, Stephan P, Dounskaia, Natalia, Walter, Charles B, & Serrien, Deborah J. (1997). Preferred and induced coordination modes during the acquisition of bimanual movements with a 2: 1 frequency ratio. Journal of Experimental Psychology: Human perception and performance, 23(4), 1087.

Team, R Core. (2018). R: A Language and Environment for Statistical Computing. Vienna, Austria: R Foundation for Statistical Computing. Retrieved from https://www.r-project.org/

Tsutsui, Seijiro, Lee, Timothy D, & Hodges, Nicola J. (1998). Contextual interference in learning new patterns of bimanual coordination. Journal of motor behavior, 30(2), 151–157.

VaezMousavi, SM, & Rostami, R. (2009). The effects of cognitive and motivational imagery on acquisition, retention and transfer of the basketball free throw. World Journal of Sport Sciences, 2(2), 129–135.

Wu, Will FW, Young, Doug E, Schandler, Steven L, Meir, Gily, Judy, Rachel LM, Perez, Jonae, & Cohen, Michael J. (2011). Contextual interference and augmented feedback: is there an additive effect for motor learning? Human movement science, 30(6), 1092–1101.

Yeganeh Doost, Maral, Orban de Xivry, Jean-Jacques, Bihin Benoît, & Vandermeeren, Yves. (2017). Two Processes in Early Bimanual Motor Skill Learning. Frontiers in human neuroscience, 11, 618–618. doi: 10.3389/fnhum.2017.00618

